# A Novel Approach on dTMS Coil Design Based on Tractography and Neural Cable Theory

**DOI:** 10.1101/683870

**Authors:** Ali Mohtadi Jafari, Ali Abdolali

**Affiliations:** BioElectromagnetic Group, Applied Electromagnetics Laboratory, School of Electrical Engineering, Iran University of Science and Technology, Tehran, Iran

**Keywords:** deep brain stimulation, reciprocity theorem, transcranial magnetic stimulation

## Abstract

Deep transcranial magnetic stimulation (dTMS) plays a useful role in the treatment of many diseases. In previous designs, it is only desired to maximize the absolute value of the electric field in the target zone. Whereas due to the nerve cable equation (NCE), the membrane potential of a nerve cell is changed due to the rate of change in the tangential electric field 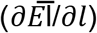. In this paper, the criterion of the design is to maximize 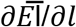 along the nerve cell fiber and simultaneously minimize the absolute value of the electric field. The direction of the nerve fibers is obtained using Tractography. Then, the specifications of source and coils such as position and spatial angles are determined in such a way that every two coils cancel the absolute value of the electric field of each other by using reciprocity theorem. This happened whereas the 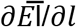 is maximized. For better comparison, a 20-coil array is designed for stimulation of cingulum region of the brain. Results of this array are compared with results of the FO8-Halo coil and it is shown that with the same value for 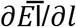, an absolute value of the induced field in the brain for the 20-coil array is much smaller than the FO8-Halo coil. Also, designed coil array requires smaller sources. So, the array can stimulate the target zone more accurately and with higher efficiency. This new technique can solve the problem of stimulation of the large areas of the brain and stimulate the desired target zone with more accuracy.

## 1) Introduction

TRANSCRANIAL MAGNETIC STIMULATION (TMS) [1] is a non-invasive technique that is widely used for brain stimulation. TMS is useful in treatment and enhancement of skills in many diseases such as Parkinson’s and depression [2]. In TMS method, a time variant magnetic field is produced in the brain. Such magnetic field leads to induction of electric field in the brain. Based on neuronal cable equation [3], the induced electric field can change the voltage of the cell membrane and so excite the nerve cell. The goal of the effort in many previous works is to excite the cortex region [4]. However, stimulation of the deeper regions of the brain has an important role in the treatment of many diseases [5].

Deep Brain Stimulation(DBS) is the surgical procedure used to stimulate the deeper regions of the brain. In DBS technique, an electrode is placed in the brain by using surgery. This electrode is connected to pacemaker-like devices and so electrical field is directly exerted to the one or multi-region of the brain [6]. Stimulation of each special region is efficient in the treatment of special diseases. For example, stimulation of internal capsules [7], nucleus accumbens [8], [9], nucleus basalis of Meynert [10], caudate nucleus [11] and Cingulum [12] is efficient in treatment of obsessive-compulsive disorder (OCD), disorder-major depression, Alzheimer, and chronic neuropathic pain.

The need of surgery and difficulty of exerting E-field in this method, are two major deficiencies that cause to effort of using TMS method for non-invasive stimulation of the deep regions too [13]. So deep transcranial magnetic stimulation (dTMS) is a new technique that requires the new coils for deeper stimulation. dTMS can effectively treat many diseases such as apathy, major depressive disorder (MDD), auditory hallucinations of schizophrenia, and bipolar depression, as well as improve cognitive [14], [15].

Nowadays several structures [16] such as H-Coil [17] and Halo Coil [18]–[20] have been used for dTMS. Coil arrays are among the most useful structures in TMS systems [21]–[23]. Coil arrays are more flexible respect to other structures because they have much more independent design parameters. However, problems exist in the way of using these structures such as a requirement of several sources [23].

The main problem in dTMS is the impossibility of having a real focus in a target zone [24]. So for stimulation of deep regions, a large volume of the brain is stimulated. Indeed, in all coils designed in previous works, always the absolute value of the electric field is considered as stimulation criterion [25]. So it is only desired to increase the absolute value of the electric field in deep regions and desired stimulation regions whereas according to the nerve cable equation [3] the changes of the tangential electric field along the nerve fiber is mainly change the voltage of the nerve cell membrane.

In recent years, the direction of nerve cells has been detected by using advanced brain imaging technique called tractography based on diffusion tensor imaging (DTI) [26]. In previous researches, it is shown that the orientation of nerve cells in the brain has a major effect in stimulation of the nerve fiber using TMS [27]–[29].

In our previous work [30], a method based on reciprocity theorem has been proposed for the design of the coil arrays. In this method, without the need of optimization algorithms, the best location and spatial angle of the coils are determined in such a way that the maximum field is induced in the desired direction (in the direction of the neural cell) at the target location. As it is known, the location and angle of coils have a great influence on the induction field at the target location. In [30], the proposed method was used to design an array with the goal of maximizing the absolute value of the tangential field (|*Ē*_*l*_|) on the neural cell.

In this paper, that reciprocity-based method is used with a small amount of variation to introduce a new design approach for stimulation systems for dTMS applications. In this work, a closer look at the nerve cell cable equations is made. In this approach, the ultimate goal is to design more effective and accurate stimulation systems based on tractography and cable equations. Accordingly, the aim is to maximize tangential field variations along the nerve cell 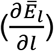 and simultaneously, contrary to past work, minimize the absolute value of the electric field (|*Ē*|) in the target region. In this paper, this approach is implemented by coil arrays. However, this approach can be used for any other stimulation systems.

This new approach leads to design coil arrays with higher efficiency and smaller sources. Also, coil arrays will be designed with respect to the real direction of nerve cells that are obtained from tractography. By obtaining the direction of nerve cells, it is being able to design coils based on tangential field variations along the nerve cell 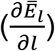.

By this approach, coil arrays will be designed in such a way that every two coils cancel the absolute value of the electric field of each other and simultaneously the rate of change in the tangential induced field will be maximized. The results of this approach are to obtain maximum stimulation in target zone whereas achieving the minimum absolute value of the electric field in the whole brain. The organization of this paper is as follows: In second section, principles of operation including reciprocity theorem and design method will be completely described. Also, an example of an array with two coils will be solved. Finally, third Section is devoted to designing of the 20-coil array for stimulation of the cingulum region of the brain. In addition, numerical results are compared with the results of FO8-Halo coils.

## 2) Materials and Methods

For the better understanding of the design approach, it will be more appropriate to begin by focusing attention on the nerve cable equation. This equation shown in (1), describes that how the potential of the nerve cell membrane is changed due to the external electric field [3].

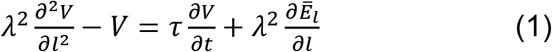

In this equation, V is membrane potential, λ is cell membrane length constant, τ is the time constant and *Ē*_*l*_ is the tangential part of the induced electric field. The schematic part of the nerve cell is shown in Figure 1a. If it is assumed that this nerve cell is along 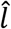, then based on (1), only the tangential electric field plays a role in stimulation of the cell. Also, it can be understood from (1) that change of the tangential electric field along the nerve fiber (i.e. 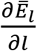) can caused change in the membrane potential.

**Figure 1.**
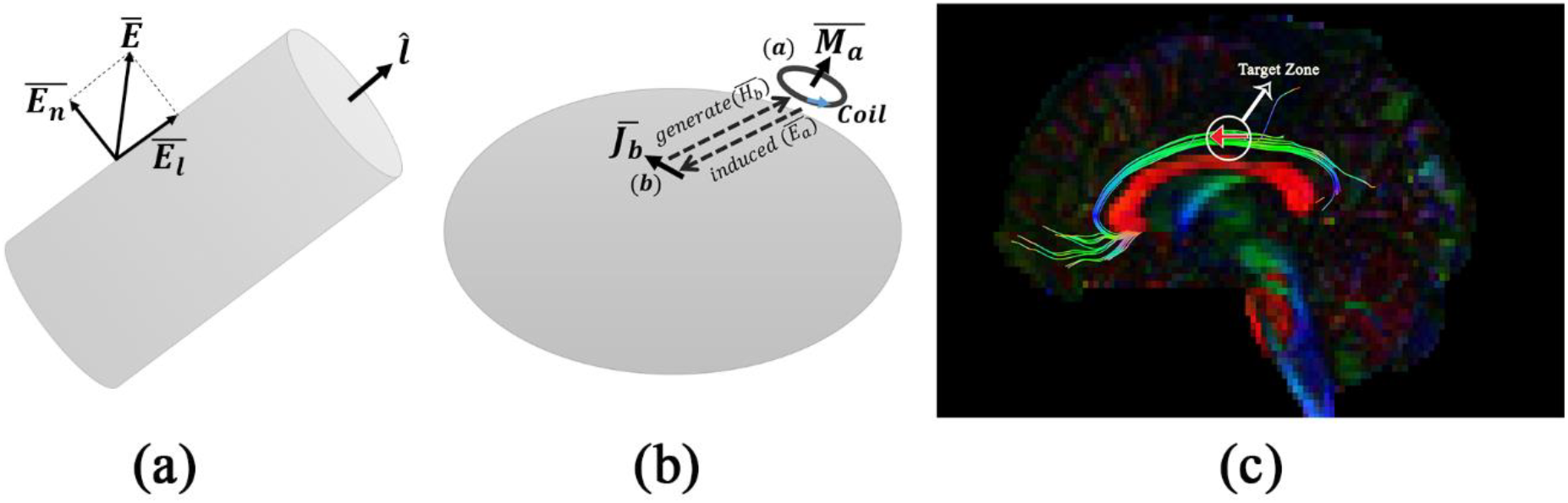
(a) Schematic crosscut of a neuron, its local direction and components of induced electric field on the surface. (b) Schematic view of the TMS problem and the way it is adopted to reciprocity theorem. (c) Target zone as a part of nerve fibers of cingulum. Tractography shown the nerve direction of target zone.

Based on (1) for designing of the efficient stimulation system, it is necessary to maximize the 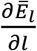. So unlike the previous works, the goal of the effort in proposed design method is to simultaneously maximize the 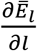 and minimize the absolute value of the electric field. With this approach, in addition to achieving the efficient stimulation system, fewer regions will be stimulated undesirably.

### Reciprocity theorem in magnetic stimulation of the brain

Reciprocity theorem is among the most important electromagnetics theorems [31]. This theory can be shown in inhomogeneous media as (2).

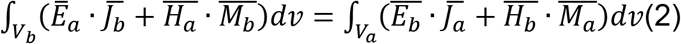

In this equation, 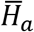 and *Ē*_*a*_ are the magnetic fields and electric fields generated by electric current 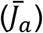 and magnetic current 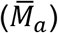 at point (b). Similarly, 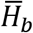 and *Ē*_*b*_ are the magnetic fields and electric fields generated by 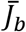 and 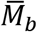 at point (a).With the help of this theory, it can be shown that position of the sources and where the induced fields are measured can be interchanged. In the other words, this theory can relate the induced field at the first source due to the second source to the induced field at the second source due to the first source [31].

For better illustration, (2) is simplified for TMS design problem and conceptually shown in Figure 1b. In TMS system, induced electric field is being applied by the one or more numbers of coils. According to the Figure 1b, it is assumed that the target region of (b) is inside the head and coil is positioned at point of (a) outside of the head.

It is known by the electromagnetic theory that a coil can be modeled as a magnetic dipole with a magnetic current of 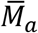[24], [32]. So if sources are assumed to be point sources, (2) will be simplified as (3).

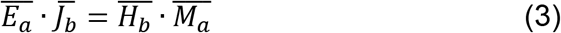

This equation determines the relation between the induced electric field of *Ē*_*a*_ by a coil with the magnetic current of 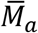 at the target point of (b) and magnetic field of 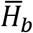 by the electric dipole with an electric current of 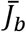 at the position of the coil (a). Hence for designing the efficient magnetic stimulation, the electric dipole must be put at the target point with direction as same as nerve fiber 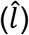. So the electric current of the dipole can be shown as (4).

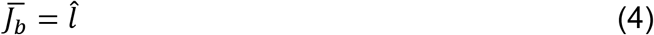

In consequence (3) transforms to (5).

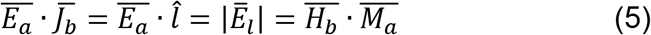

Now it is sufficient to calculate the magnetic field generated by the electric dipole at the point of (a) with any desired method such as numerical methods. After that, the value of the effective induced electric field (i.e. *Ē*_*l*_) can be obtained and coil array can be designed based on the following roles:

1. Due to the dot product between 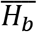 and 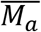 in (5), for maximization of *E*_*l*_, 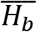 must be parallel to the 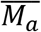. Based on this rule, the optimum spatial angle of each coil is determined.
2. The bigger the value of 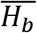 at the position of a coil is, the bigger the *Ē*_*l*_ induced by the similar coil will be. Hence the optimum position of the coils is determined based on this rule.
3. If 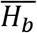 has similar values at the position of two similar coils with same sources, these two coils will have similar induced *E*_*l*_ that cause to cancels each other.

These rules help a user to determine the position and spatial angle of each coil of an array and simultaneously give a user a proper attitude about the interaction between the coils of an array in induction of electric field in order to determine the coils that cancel each other.

### Design approach

This subsection is devoted to designing of the two-element coil array. Target zone is assumed to be a part of nerve fibers of cingulum section of the brain as shown in Figure 1c. ExploreDTI software is used to determine the direction of nerve cells in this area [33]. Also for stimulation of the head, the real HUGO model of the head and the low-frequency solver of the CST software are used [34]. Figure 2 shows designed array above the head.

**Figure 2.**
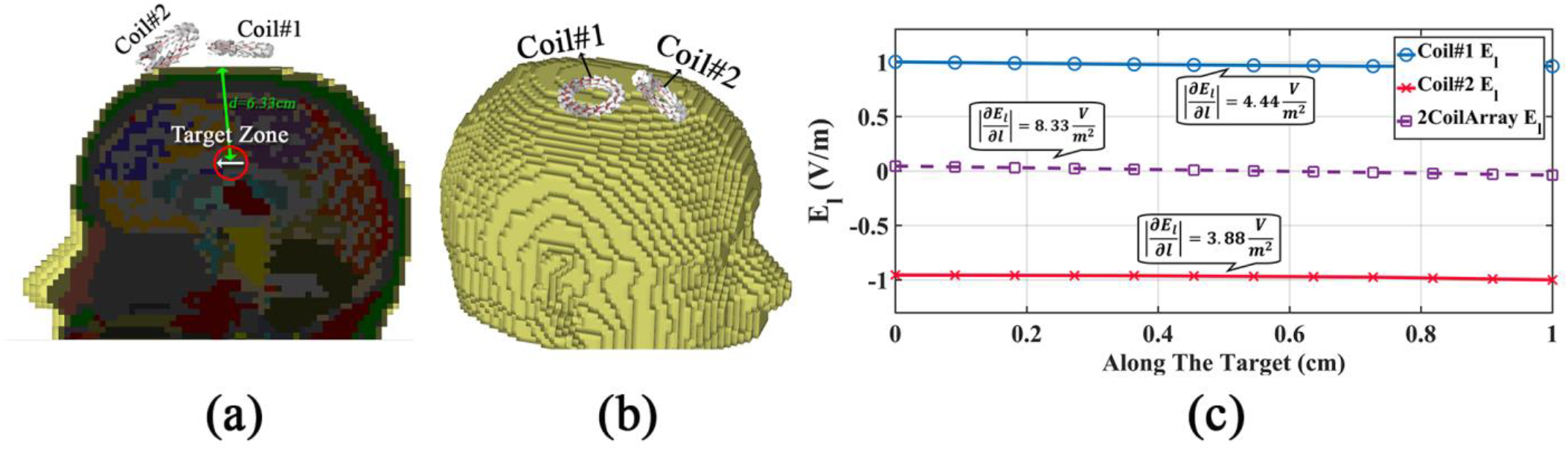
(a) Cross cut of HUGO model at the plane of Target zone. (b) 2-Coil array around HUGO model. (c) Tangential component of induced Electric field from each coil and 2coil array and its 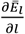 on target zone.

According to the Figure 2a at the first step, an electric dipole with the same direction as nerve cells is placed in the target zone. After solving the problem using low-frequency solver of CST software, the induced magnetic field outside the head is calculated and then the position of coils is determined. Based on calculated 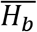, the two-coil array is designed and shown in Figure 2b. the spatial angle of both coils is determined based on the direction of 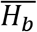.

An important note in this design is that 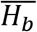 in the position of these two coils has the same amplitude and this means that if these coils have sources with same absolute values and inverse directions (opposite sign), they will cancel the induced field of each other at the target zone. There is no need of two sources in this case. It is sufficient to inverse the wiring direction of coils and connects both of them to a source. In fact these two coils have the same sources and opposite current directions.

The 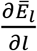 generated along the nerve fiber is shown in Figure 2c. It can be seen that however these coils cancel induced electric field of each other, 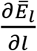 of them is effectively gathered together. So based on (1) the nerve fibers of the target zone can be stimulated. As already mentioned, the goal of the design is to minimize the electric field, including both vertical and tangential components. However, since, with the help of the reciprocity theorem, angle and location of the coils are determined in such a way that the majority of the induction field is tangential, the effect of the vertical component is ignored in the design algorithm.

## 3) RESULTS

In the previous section, the design aspects of coil arrays were indicated and an example of the two-coil array was designed for cingulum section of the brain. This section is devoted to the design of the 20-element coil array for stimulation of the region shown in Figure 3. Positions of coils and distance between the array and the head are determined in such a way that each coil can be rotated freely without any collision with the head and other coils.

**Figure 3.**
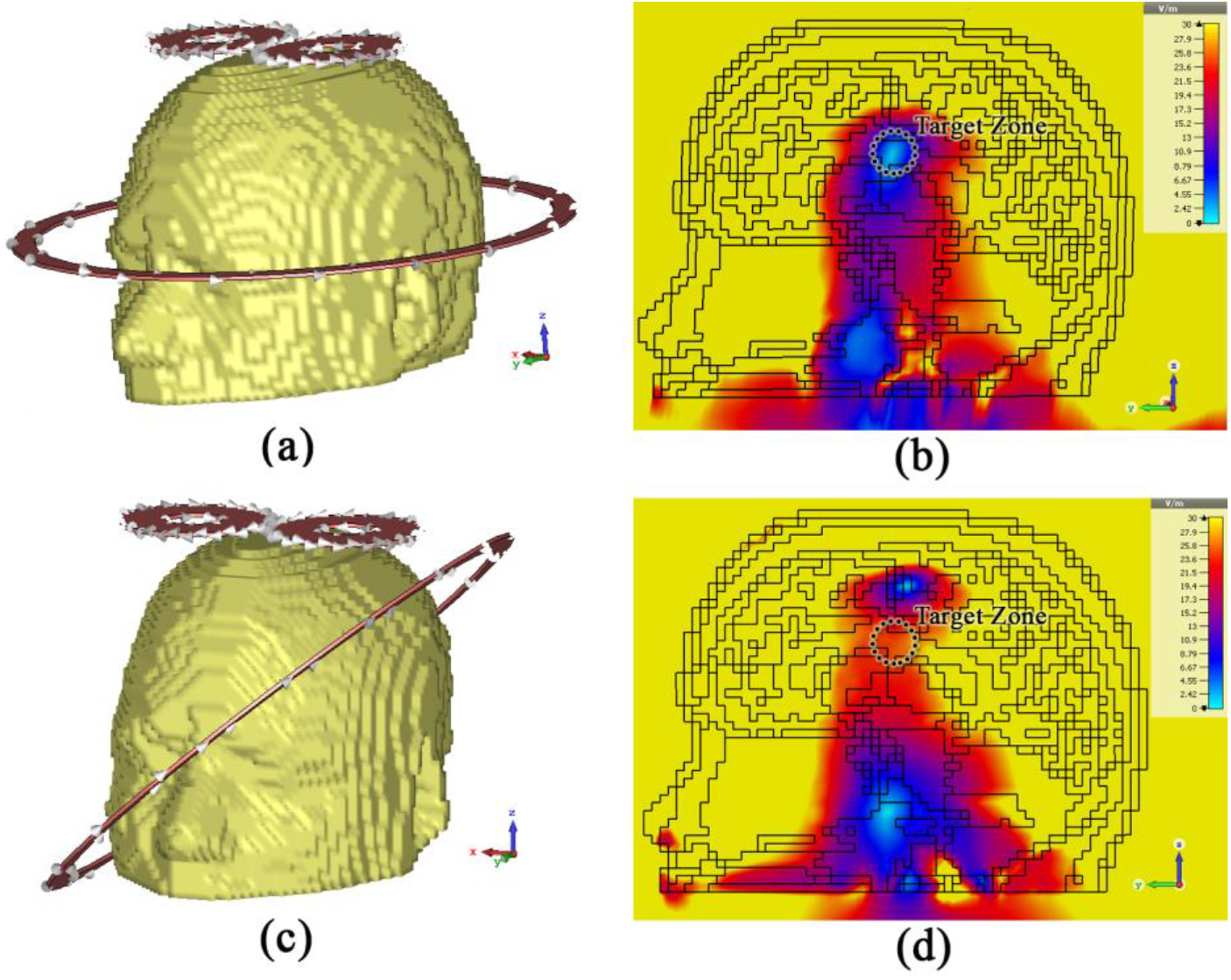
(a) Halo-FO8 Coil and Hugo model. (b) Induced electric filed (|*Ē*|) in brain from Halo-FO8 Coil. (c) 40° rotated Halo-FO8 coil around HUGO model (d) Induced electric field (|*Ē*|) in brain from 40° rotated Halo-FO8 coil.

As same as a two-coil array, for each two coils of this array, absolute values of the induced field of *Ē*_*l*_ are canceled each other whereas 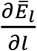 of coils gather together.

For better comparison, let’s consider a Halo-FO8 coil as it is shown in Figure 3a besides the HUGO model. Specifications of this coil are also described in Table 1.The induced electric field of the FO8-Halo coil in the whole brain is shown in Figure 3b. Halo coil was first presented for dTMS applications in 2011 [18]. In 2015 [19] performance of this coil was improved by upgrading to the FO8-Halo coil. Finally, variable “Halo coil” was introduced in 2015 [20] with the ability to rotate its larger coil for stimulation of different regions of the brain.

**TABLE 1.** HALO-FO8 COIL PARAMETERS^29^

| Coils Type | N*I<br>(TURN* CURRENT (A)) | $R_{in}(cm)$ | $R_{out}(cm)$ |
| --- | --- | --- | --- |
| FO8 | 50000 | 1.5 | 4 |
| Halo | 25000 | 13.8 | 15 |

In this case, for maximization of stimulation in the target zone, it is assumed that larger coil is rotated about 40 degrees as shown in Figure 3c. Also, induced electric field of the FO8-Halo coil in the brain for 40 degrees rotated case is shown in Figure 3d. For the better comparison, values of the induced electric field along the target nerve fiber with and without rotation are shown in Figure 4. As it can be seen, with rotation better stimulation can be obtained. In Figure 4 for rotated Halo coil, the rate of change in the tangential induced field of *Ē*_*l*_ is as (6).

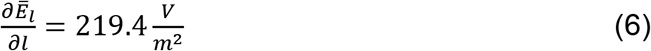

**Figure 4.**
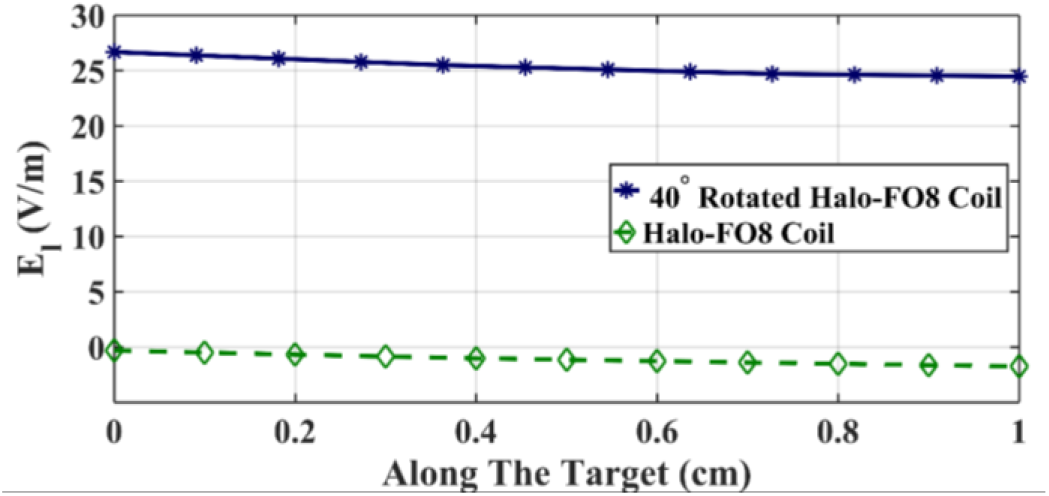
Induced tangential electric field from Halo-FO8 Coil and 40° rotated Halo-FO8 coil in target zone.

As can be seen from Figure 3b and Figure 3d, for stimulating the nerve fibers of the cingulum section almost whole brain is under the undesired electric field. Now, Based on the value of 219.4 v/m^2^ for 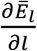, a coil array with 20 coils is designed.

The specifications of coils are described in Table 2 and is shown in Figure 5a. Also, induced electric field and tangential electric field of *Ē*_*l*_ are shown in Figure 5b and Figure 5c.

**TABLE 2.** 20-COIL ARRAY PARAMETER

| Coils Type | N*I<br>(TURN* CURRENT (A)) | $R_{in}(cm)$ | $R_{out}(cm)$ |
| --- | --- | --- | --- |
| 20-Coil<br>Array | 20200 | 1.5 | 2 |

**Figure 5.**
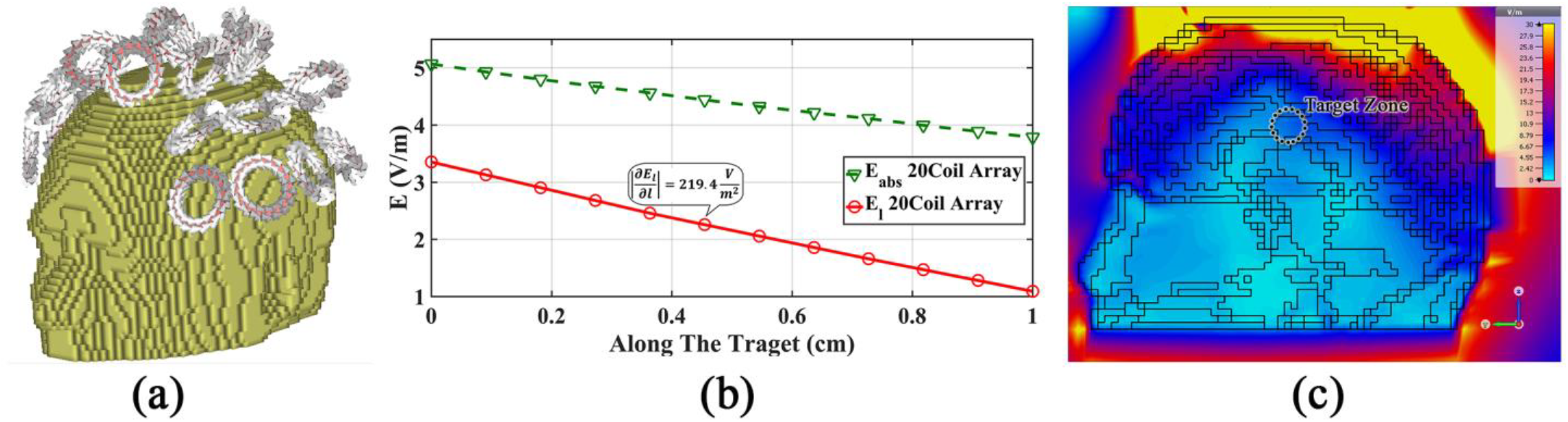
**(a)** Designed 20-coil array around the head. **(b)** Induced electric field and its tangential component from designed 20 coil array along target zone. **(c)** Induced electric field (|*Ē*|) in brain from designed 20-coil array.

## 4) DISCUSSION

The main idea of this work began with the lack of focus in dTMS applications. As can be seen in Figure 3b and Figure 3d, this fact causes the large volume of the brain to be affected by the high intensity of the electric field in the previously designed systems. This happened because based on (1) and Figure 1a, what changes the cell membrane potential and ultimately stimulates the nerve cell is the changes of the tangential component of the electric field along the nerve fiber. With the advancement of imaging techniques and with the help of tractography, it is possible to obtain the direction of the nerve cells in the brain as shown in Figure 1c.

These three facts have raised the question of whether it is possible to minimize the intensity of the electric field in the target region and other regions of the brain, while the changes of the tangential field along the nerve fiber maximize.

To answer this question, part of the cingulum fiber was considered. This region was chosen because of the fact that, in addition to placement of this region in deep areas of the brain, tractography and its precise location in the electromagnetic model of the brain (HUGO model) were available. For initial realization, a two-coil array was considered as shown in Figure 2. These two coils were chosen based on reciprocity theorem and excited by a single current source, which neutralized the effect of the tangential electric field, while, according to Figure 2c, increase the 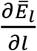 of each other. After proof of the idea, a 20-coil system was designed with the same approach.

For comparison of performance, the FO8-Halo coil was considered as Figure 3. The 20-coil system was designed in such a way that reaches the same 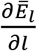 as the FO8-Halo coil. This means that, based on (1), the same performance as FO8-Halo coil in the target area must be achieved.

The result of the design is shown in Table 2. Figure 5 show the performance of this system. According to Figure 5b and (1), FO8-Halo Coil and 20-coil array have the same performance by about 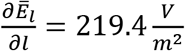. Whereas, by comparing of Figure 5c and Figure 3d, much less volume of the brain was undesirably affected by high-intensity electric field. However based on what has been said and (1), to find the points under real stimulation it is required to calculate the 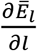 in all regions of the brain, less electric field size can at least prevent unwanted stimuli in the curves of the neural fibers.

In addition, based on table 1 and table 2, the 20-coil array designed with the proposed approach requires less current source than the FO8-Halo Coil for the same function in the target area.

In fact, based on this approach, specific designs can be made for certain regions of the brain in such a way that the stimulation and required 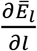 are achieved and simultaneously stimulation procedure need of smaller resources and causes to less electric field size in all areas of the brain.

It is true that real focus cannot be achieved, but this approach, while not causing the focus, can be an alternative to the lack of focusing and the inefficient performance of dTMS systems. Design based on this approach and the placement criteria 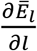 instead of the size of the electric field (|*Ē*|) can create a fundamental change in the design of magnetic stimulation systems and lead to a new generation of these systems.

This approach can be applied not only to dTMS systems but also to TMS systems. The advancement of tractography methods and achieving a more accurate map of the direction of the neurons can lead to improving the design procedures based on this approach. Future work could include dedicated designs for critical brain areas in dTMS applications.

